## Supporting information for "Anionic phospholipids stimulate the proton pumping activity of the plant plasma membrane P-type H^+^-ATPase"

This file includes all supplementary information for the manuscript:

Supplemental Figures S1, S2 and S3

Supplemental Tables S1, S2, S3, S4 and S5

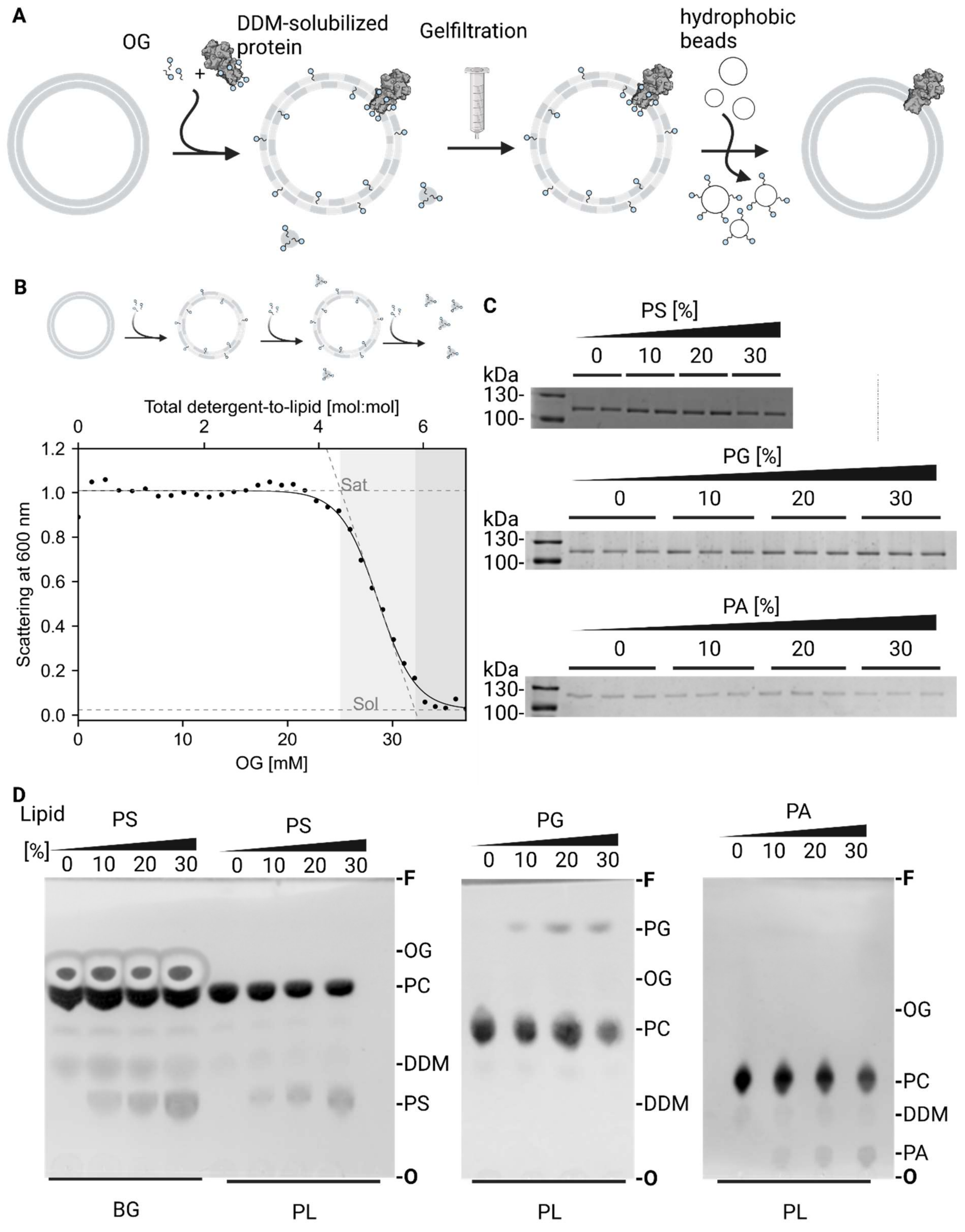


***Supplemental Figure S1. Characterization of AHA2 proteoliposomes by thin-layer chromatography and SDS-PAGE.*** *(A) Schematic diagram illustrating the reconstitution of AHA2. Preformed liposomes composed of PC in mixture with the indicated amounts of anionic phospholipids are detergent-destabilized and mixed with the detergent-solubilized $H^+$-ATPase (I). Subsequent removal of the detergent by Sephadex G-50 gel filtration (II) and Bio-Bead treatment result in the formation of sealed proteoliposomes (III). (B) Liposome solubilization is shown schematically (upper panel): liposomes (I) are destabilized upon detergent addition until completely saturated with detergent (II).*

*Further additions lead to partly solubilization and occurrence of lipid-detergent micelles (III) until liposomes are complete solubilized (IV). The procedure can be followed by light scattering at 600 nm (lower panel) as solubilization of LUVs is accompanied by less light scattering. The obtained signal was fitted to a Boltzmann equation (y=Max+(Max-Min)/(1+(1/e^(x-V50)/slope) to derive characteristic parameters such as the turning point, which was used for reconstitution and was usually slightly above the cmc of used detergent OG. (C) Coomassie Brilliant Blue-stained SDS-PAGE on AHA2 proteoliposomes, demonstrating equal amounts of protein reconstituted. Molecular weight markers are indicated on the left. Experiments were performed in duplicate (PS) or triplicate (PG, PA). (D) Liposome composition analysis before and after reconstitution using thin layer chromatography. Chromatograms shown were dried completely before photographing after staining with primuline. Difference in Rf values derive from varying temperature and running chamber sizes. OG and DDM was compared to a standard spot of 0.5 mM allowing concluding that less than 0.5 mM of each detergent remained in the sample. Different lipid percentage were analyzed by brightness of the detected spots and dividing the varying phospholipid PS, PA or PG by the sum of the test lipid and the basic lipid PC. O, origin; F, solvent front, of the chromatograms. DDM, N-dodecyl-β-maltoside; OG, octyl glucoside; PC, phosphatidylcholine; PS, phosphatidylserine; PA, phosphatidic acid.*

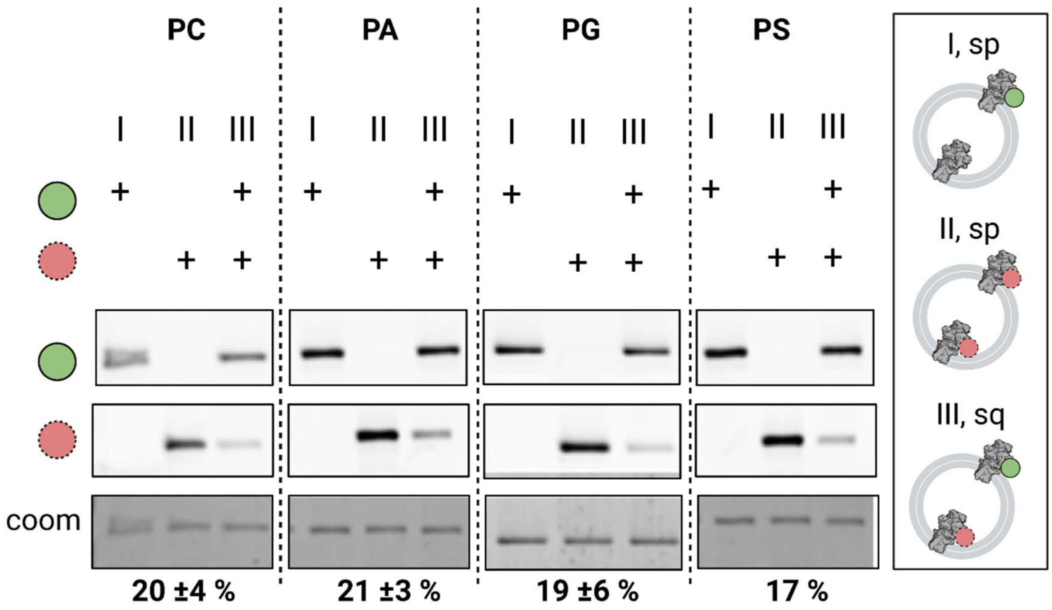


***Supplemental Figure S2. Membrane orientation of reconstituted AHA2.*** *The orientation of AHA2 in the different lipid compositions was determined by site-specific fluorescent labelling of the SNAP tag on either outward or inward facing AHA2 (ATPase domain facing outward or inward). Vesicles were incubated separately (sp) or sequentially (sq) with membrane impermeable (mi) and membrane permeable (mp) SNAP dyes and analysed as described in Material and Methods. Representative fluorescent scans (mi, mp) and corresponding Coomassie Brilliant Blue (co) stained SDS-PAGE for the indicated proteoliposome preparations (10 mol% test lipid), showing orientations of reconstituted AHA2. Numbers under the gel refer to the percentage of inward facing AHA2 (ATPase domain facing inward), determined from at least two independent reconstitutions except for PS.*

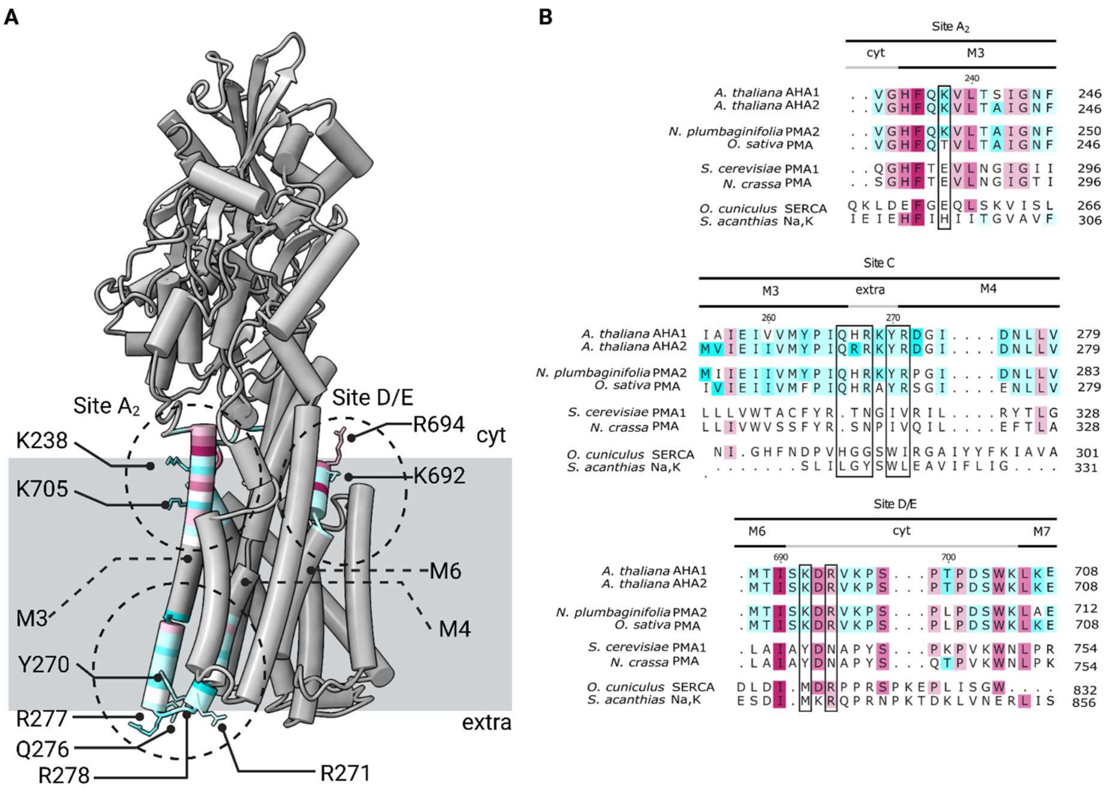


**Supplemental Figure S3*. Location of predicted phospholipid contact sites B, C, D and E in AHA2.*** (A) Cartoon representation of the AHA2 membrane domain. Patches of apprx. 15 residues around the assigned contact residues are colored according to their sequence conservation based on AL2CO algorithm implemented into ChimeraX (70). Preferentially interacting residues identified in the lipid-protein contact analysis are shown in atom representation. The approximate regions of the anionic lipid contacts sites, as identified in the lipid density maps, are marked with black, dashed circles. (B) Sequence alignment showing the conservation of residues based on AL2CO algorithm in selected P-type ATPase proton pumps from plants and yeasts as well as SERCA and Na-K-ATPase at the lipid contact sites $A_2$, C, D and E identified in AHA2. The sequences are obtained from the Uniprot database: *Arabidopsis thaliana* $H^+$-ATPase isoforms 1 and 2 (P20649 and P19456, respectively), shown in the first block, followed by the plant transporters *Nicotiana plumbaginifolia* $H^+$-ATPase isoform 2 *(Q42932), Oryza sativa subsp. Japonica* $H^+$-ATPase *(Q7XPY2),* and the two fungal transporters (*Saccharomyces cerevisiae* $H^+$-ATPase isoform 1 (P05030) and *Neurospora crassa* $H^+$-ATPase (P07038). Lastly, they are compared to the SERCA pump from rabbit *Oryctolagus cuniculus* (P04191) and the Na,K-ATPase from shark *Squalus acanthias* (Q4H132). Residues in black boxes are found to have enriched lipid contacts in the MD simulations.

**Supplemental Table S1:** Statistical analysis of the data presented in Figure 1. Mean of indicated measurements is displayed together with standard deviation. Data was undertaken pairwise t-test and obtained p-values were group – p > 0.05: not significant (ns), p < 0.05: significant (*), p < 0.01: very significant (**), p < 0.001: highly significant (***)

| | count | mean | std | PC | PE | PA | PG | PS | Mix | group |
|---|---|---|---|---|---|---|---|---|---|---|
| **PC** | 12 | 15.4 | 8.1 | - | ns | ns | * | *** | ** | b |
| **PE** | 4 | 10.4 | 6.8 | | - | ns | ns | *** | ** | b |
| **PA** | 3 | 34.0 | 8.6 | | | - | ns | *** | ns | abc |
| **PG** | 7 | 36.9 | 8.9 | | | | - | *** | ns | ac |
| **PS** | 9 | 83.9 | 25.1 | | | | | - | ** | d |
| **Mix** | 5 | 49.2 | 16.0 | | | | | | - | a |

***Supplemental Table S2:** Statistical analysis of the data presented in Figure 2A (PA titration). Mean of indicated measurements is displayed together with standard deviation. Data was undertaken pairwise t-test and obtained p-values were group – p > 0.05: not significant (ns), p < 0.05: significant (*), p < 0.01: very significant (**), p < 0.001: highly significant (***)*

| | count | mean | std | 0 mol% | 10 mol% | 20 mol% | 30 mol% | group |
|---|---|---|---|---|---|---|---|---|
| **0 mol%** | 12 | 15.4 | 8.1 | - | ns | ** | ns | a |
| **10 mol%** | 4 | 27.6 | 17.3 | | - | ns | ns | ab |
| **20 mol%** | 2 | 50.3 | 23.2 | | | - | ns | b |
| **30 mol%** | 3 | 34.0 | 8.6 | | | | - | ab |

***Supplemental Table S3:** Statistical analysis of the data presented in Figure 2B (PG titration). Mean of indicated measurements is displayed together with standard deviation. Data was undertaken pairwise t-test and obtained p-values were group – p > 0.05: not significant (ns), p < 0.05: significant (*), p < 0.01: very significant (**), p < 0.001: highly significant (***)*

| | count | mean | std | 0 mol% | 10 mol% | 20 mol% | 30 mol% | group |
|---|---|---|---|---|---|---|---|---|
| **0 mol%** | 12 | 15.4 | 8.1 | - | ns | ** | ** | a |
| **10 mol%** | 2 | 28.3 | 21.1 | | - | ns | ns | ab |
| **20 mol%** | 3 | 42.4 | 16.5 | | | - | ns | b |
| **30 mol%** | 7 | 36.9 | 8.9 | | | | - | b |

***Supplemental Table S4:** Statistical analysis of the data presented in Figure 2C (PS titration). Mean of indicated measurements is displayed together with standard deviation. Data was undertaken pairwise t-test and obtained p-values were group – p > 0.05: not significant (ns), p < 0.05: significant (*), p < 0.01: very significant (**), p < 0.001: highly significant (***)*

| | count | mean | std | 0 mol% | 10 mol% | 20 mol% | 30 mol% | group |
|---|---|---|---|---|---|---|---|---|
| **0 mol%** | 12 | 15.4 | 8.1 | - | ns | *** | *** | b |
| **10 mol%** | 4 | 32.1 | 16.2 | | - | * | *** | b |
| **20 mol%** | 3 | 68.6 | 14.1 | | | - | ns | a |
| **30 mol%** | 9 | 83.9 | 25.1 | | | | - | a |

**Supplemental *Table S5. Percentage of time protein residues are in contact with lipid headgroups.*** *For PS headgroups, results are given for all three independent repeats and their average. For neutral headgroups, only the average is given. The tables are sorted by average PS contacts, including only residues with ≥5% PS contacts on average. Residues which show no specificity towards PS lipids are greyed out, with a ratio of average neutral:PS contacts of 80:20 or less; 90:10 is the lipid mixture.*

**PC:PS (90:10)**

| Name | ID | PS | | | | neutral |
|---|---|---|---|---|---|---|
| | | run 1 | run 2 | run 3 | Av. | Av. |
| Arg | 271 | 28 | 26 | 29 | 27 | 71 |
| Tyr | 270 | 26 | 26 | 27 | 26 | 42 |
| Lys | 707 | 21 | 25 | 32 | 26 | 74 |
| Arg | 267 | 21 | 26 | 27 | 24 | 60 |
| Lys | 60 | 30 | 17 | 21 | 23 | 77 |
| Arg | 268 | 26 | 19 | 22 | 22 | 58 |
| Lys | 57 | 28 | 15 | 19 | 21 | 39 |
| Phe | 61 | 25 | 14 | 21 | 20 | 60 |
| Lys | 692 | 19 | 20 | 18 | 19 | 79 |
| Lys | 705 | 14 | 21 | 22 | 19 | 38 |
| Lys | 238 | 21 | 12 | 23 | 19 | 34 |
| Trp | 667 | 19 | 19 | 19 | 19 | 66 |
| Lys | 733 | 17 | 29 | 10 | 19 | 62 |
| Trp | 66 | 22 | 13 | 19 | 18 | 66 |
| Arg | 842 | 11 | 20 | 22 | 18 | 61 |
| Leu | 706 | 19 | 12 | 20 | 17 | 28 |
| Arg | 694 | 15 | 17 | 19 | 17 | 63 |
| Gln | 266 | 17 | 15 | 17 | 17 | 79 |
| Asn | 67 | 12 | 12 | 15 | 13 | 76 |
| Thr | 241 | 12 | 10 | 18 | 13 | 57 |
| Asn | 245 | 13 | 9 | 17 | 13 | 64 |
| Thr | 734 | 12 | 21 | 5 | 13 | 76 |
| Tyr | 804 | 12 | 10 | 16 | 13 | 75 |
| Gly | 816 | 9 | 12 | 15 | 12 | 46 |
| Trp | 817 | 9 | 12 | 14 | 12 | 69 |
| Ile | 666 | 12 | 14 | 10 | 12 | 40 |
| Val | 743 | 8 | 14 | 12 | 12 | 73 |
| Lys | 269 | 14 | 9 | 11 | 11 | 18 |
| Ala | 785 | 10 | 11 | 13 | 11 | 47 |
| Arg | 89 | 9 | 9 | 16 | 11 | 88 |
| Asn | 85 | 11 | 8 | 14 | 11 | 76 |
| Thr | 689 | 10 | 10 | 12 | 11 | 45 |
| Glu | 708 | 5 | 13 | 13 | 11 | 33 |
| Ser | 691 | 11 | 11 | 10 | 11 | 63 |
| Leu | 786 | 10 | 11 | 10 | 10 | 35 |
| Phe | 710 | 11 | 8 | 11 | 10 | 27 |
| Trp | 729 | 9 | 15 | 6 | 10 | 39 |
| Asn | 806 | 12 | 4 | 13 | 10 | 87 |
| Phe | 64 | 14 | 6 | 8 | 9 | 23 |
| Gly | 784 | 8 | 9 | 12 | 9 | 42 |
| Pro | 783 | 8 | 10 | 10 | 9 | 69 |
| Arg | 744 | 7 | 11 | 10 | 9 | 26 |
| Met | 688 | 9 | 10 | 9 | 9 | 58 |
| Leu | 59 | 11 | 9 | 8 | 9 | 42 |
| Asn | 115 | 10 | 10 | 6 | 9 | 69 |
| Ile | 265 | 8 | 9 | 9 | 9 | 40 |
| Val | 803 | 7 | 8 | 11 | 9 | 40 |
| Ala | 242 | 8 | 6 | 11 | 9 | 19 |
| Phe | 111 | 12 | 8 | 5 | 8 | 45 |
| Gly | 818 | 6 | 11 | 8 | 8 | 40 |
| Asn | 116 | 10 | 6 | 9 | 8 | 69 |
| Tyr | 843 | 6 | 9 | 8 | 7 | 83 |
| Phe | 736 | 7 | 14 | 1 | 7 | 72 |
| Phe | 669 | 7 | 11 | 4 | 7 | 18 |
| Thr | 740 | 7 | 11 | 3 | 7 | 73 |
| Trp | 71 | 6 | 6 | 9 | 7 | 46 |
| Phe | 741 | 6 | 8 | 6 | 7 | 74 |
| Ser | 776 | 6 | 9 | 5 | 6 | 73 |
| Gly | 86 | 6 | 6 | 7 | 6 | 20 |
| Asp | 735 | 5 | 12 | 1 | 6 | 70 |

**PC:PE:PS (45:45:10)**

| Name | ID | PS | | | | neutral |
|---|---|---|---|---|---|---|
| | | run 1 | run 2 | run 3 | Av. | Av. |
| Lys | 707 | 29 | 28 | 27 | 28 | 71 |
| Arg | 842 | 21 | 22 | 26 | 23 | 58 |
| Lys | 705 | 22 | 23 | 20 | 22 | 41 |
| Lys | 238 | 20 | 23 | 18 | 20 | 43 |
| Arg | 271 | 26 | 14 | 14 | 18 | 80 |
| Arg | 694 | 12 | 18 | 21 | 17 | 63 |
| Lys | 733 | 19 | 12 | 20 | 17 | 60 |
| Leu | 706 | 18 | 19 | 13 | 17 | 29 |
| Lys | 60 | 21 | 17 | 9 | 16 | 75 |
| Arg | 268 | 21 | 9 | 18 | 16 | 55 |
| Arg | 267 | 22 | 12 | 13 | 16 | 75 |
| Trp | 667 | 15 | 11 | 18 | 14 | 71 |
| Gly | 816 | 13 | 11 | 16 | 14 | 46 |
| Phe | 61 | 17 | 16 | 6 | 13 | 67 |
| Thr | 241 | 12 | 11 | 16 | 13 | 62 |
| Thr | 734 | 14 | 10 | 13 | 13 | 77 |
| Gln | 266 | 15 | 9 | 14 | 13 | 83 |
| Lys | 57 | 14 | 21 | 3 | 13 | 35 |
| Tyr | 270 | 23 | 7 | 7 | 12 | 52 |
| Pro | 783 | 11 | 9 | 14 | 12 | 63 |
| Trp | 66 | 15 | 13 | 6 | 12 | 68 |
| Trp | 817 | 11 | 9 | 13 | 11 | 70 |
| Glu | 708 | 10 | 10 | 13 | 11 | 29 |
| Asn | 245 | 10 | 11 | 13 | 11 | 58 |
| Asn | 85 | 11 | 12 | 11 | 11 | 77 |
| Arg | 89 | 8 | 14 | 10 | 11 | 88 |
| Arg | 782 | 13 | 8 | 11 | 11 | 52 |
| Trp | 729 | 12 | 7 | 12 | 10 | 42 |
| Ala | 785 | 6 | 12 | 12 | 10 | 48 |
| Arg | 744 | 9 | 7 | 13 | 10 | 25 |
| Val | 743 | 11 | 7 | 12 | 10 | 75 |
| Val | 803 | 10 | 8 | 11 | 10 | 39 |
| Tyr | 804 | 11 | 8 | 10 | 10 | 78 |
| Ile | 666 | 10 | 6 | 12 | 9 | 41 |
| Asn | 67 | 10 | 9 | 9 | 9 | 78 |
| Gly | 784 | 6 | 10 | 11 | 9 | 44 |
| Lys | 692 | 11 | 8 | 7 | 9 | 89 |
| Phe | 710 | 10 | 9 | 7 | 8 | 25 |
| Ala | 242 | 9 | 8 | 7 | 8 | 20 |
| Glu | 781 | 8 | 7 | 9 | 8 | 81 |
| Leu | 59 | 10 | 7 | 7 | 8 | 51 |
| Asn | 115 | 9 | 9 | 5 | 8 | 67 |
| Asn | 116 | 10 | 6 | 6 | 7 | 70 |
| Tyr | 843 | 6 | 7 | 9 | 7 | 83 |
| Ile | 265 | 10 | 5 | 6 | 7 | 50 |
| Ser | 776 | 5 | 8 | 8 | 7 | 72 |
| Phe | 839 | 5 | 7 | 8 | 7 | 60 |
| Gly | 818 | 6 | 5 | 9 | 7 | 36 |
| His | 235 | 7 | 8 | 4 | 6 | 4 |
| Phe | 741 | 5 | 6 | 6 | 6 | 77 |
| Gly | 742 | 5 | 6 | 6 | 6 | 67 |
| Val | 780 | 6 | 5 | 5 | 6 | 81 |
| Gly | 814 | 5 | 5 | 7 | 5 | 11 |
| Asn | 806 | 7 | 4 | 5 | 5 | 91 |
| Phe | 669 | 6 | 2 | 8 | 5 | 16 |
| Trp | 71 | 7 | 4 | 6 | 5 | 45 |
| Leu | 786 | 5 | 6 | 5 | 5 | 44 |
| Ile | 815 | 5 | 5 | 6 | 5 | 11 |
| Ile | 844 | 5 | 5 | 6 | 5 | 78 |
| Gly | 86 | 7 | 4 | 5 | 5 | 26 |

| | | | | | | |
|---|---|---|---|---|---|---|
| Phe | 839 | 5 | 9 | 5 | 6 | 61 |
| Met | 65 | 8 | 3 | 7 | 6 | 17 |
| Gly | 742 | 4 | 8 | 5 | 6 | 61 |
| Ala | 805 | 7 | 3 | 8 | 6 | 33 |
| Arg | 813 | 6 | 3 | 8 | 6 | 15 |
| Glu | 781 | 6 | 8 | 4 | 6 | 82 |
| Arg | 782 | 5 | 7 | 5 | 6 | 44 |
| Asp | 693 | 5 | 6 | 5 | 6 | 13 |
| Asp | 739 | 5 | 9 | 2 | 5 | 86 |
| His | 235 | 6 | 4 | 6 | 5 | 4 |

| | | | | | | |
|---|---|---|---|---|---|---|
| Trp | 93 | 3 | 7 | 5 | 5 | 42 |
